# High-LET Particle Therapy Targets Tumor Microtube Networks in Glioblastoma

**DOI:** 10.1101/2025.01.31.635916

**Authors:** Ivana Dokic, Federica Ciamarone, Dirk C. Hoffmann, Jovana Bojcevski, Damir Krunic, Thomas Tessonnier, Jürgen Debus, Andrea Mairani, Varun Venkataramani, Amir Abdollahi

## Abstract

**Purpose:** Tumor cell networks formed by tumor microtubes (TMs) are thought to drive therapy resistance in glioblastoma (GB). X-ray irradiation enhances TM formation, thereby increasing radioresistance. We hypothesize that high linear energy transfer (LET) particle radiotherapy is less affected by TM-mediated resistance due to its reduced reliance on indirect DNA damage. This study explores the impact of LET-induced DNA damage on TMs formation and GB survival

**Material and Methods:** Formation of TMs was investigated in the primary patient derived glioblastoma stem-like cell lines (S24 and T269) irradiated with different LET, ranging from 3 – 107 keV/µm, across dose series (1, 2, 4, 6 Gy) of clinical proton, helium, and carbon ion beams. TM networks and DNA damage patterns, specifically γH2AX foci, were visualized using fluorescence microscopy. Cell survival was evaluated through clonogenic survival assays.

**Results:** The formation of TMs, radiation-induced nuclear DNA damage repair foci, and GB cell survival were correlated with a gradual increase in LET. Consistent with conventional photon/X-rays, low-LET proton irradiation promoted TMs formation in a dose-dependent manner. In contrast, an anti-correlation between LET and TMs induction was found, i.e., a decreased network connectivity with gradual increase of LET and formation of complex DNA damage. Consequently, LET increase correlated with reduced cell survival, with the most pronounced cell killing observed after high-LET carbon irradiation. Moreover, the inverse correlation between LET and TMs density was further confirmed for a broad range of LET modulated within the carbon ion irradiation.

**Conclusion:** This is the first report on the relevance of LET as a novel mean to overcome TMs network-mediated radioresistance in GB, with ramifications for the clinical translation of high-LET particle radiotherapy to further improve outcome in this still devastating disease.

## Introduction

Therapy resistance in glioblastoma (GB) remains a significant challenge in its effective management [1]. Emerging research underscores the pivotal role of tumor microtubes (TMs) - elongated membrane tubes forming functional tumor cell networks - in fostering resistance to chemotherapy [2] and radiotherapy [3, 4]. It has been observed that tumor cells, when linked through TMs, display heightened endurance against the adverse impacts of conventional radiotherapy [4]. Furthermore, TMs have been recognized as integral elements in the neuron-glioma synapse and the stem-like compartment of astrocytic gliomas, thereby playing a crucial role in tumor progression [5-8].

Precision particle radiotherapy with protons, helium and carbon ions has emerged as a clinical approach with preferred biophysical characteristics and therefore greater potential for tumor treatment compared to conventional photon/X-ray irradiation techniques [9]. This is due to the Bragg peak phenomenon inherent to ion irradiation, enabling targeted delivery of the particle beam to the tumor, while minimizing collateral damage to surrounding healthy tissue [10]. Each particle modality is characterized by specific physical properties (depth-dose and lateral scattering) and particle ionization along its traversal – the Linear Energy Transfer (LET in keV/µm). These characteristics can lead to different relative biological effectiveness (RBE), representing a radiation type’s ability to cause biological damage compared to a reference, typically photon irradiation [9]. Particles generate a denser ionization trail than photons, causing more complex DNA damage even at the same physical radiation dose, particularly with high LET radiation [11]. Both low and high LET irradiation can induce DNA damage through direct and indirect mechanisms [12]. Low LET radiation predominantly causes indirect damage by ionizing surrounding molecules, like water, creating reactive oxygen species (ROS) that subsequently damage the DNA [13]. High LET radiation is more likely to cause direct DNA damage due to its dense ionization tracks that directly impact the DNA molecules inducing complex lesions [14]. Increase in complex, unrepairable and persistent DNA damage, [15, 16] strongly correlates with decrease in cell survival [16], and represents a hallmark of radiation induced anticancer effects. Complex DNA damage involves multiple types of lesions occurring in close proximity [17], and it can be visualized using γH2AX as a marker for radiation induced foci (RIF) [18].

Considering the potential of intercellular micro- and nanotubes to facilitate cell to cell communication and support ROS signaling and transfer, TMs network may play a more significant role in indirect DNA damage induced by low LET, than in direct high-LET induced-DNA damage.

In this work, TMs network induced radioresistance was addressed by studying the network formation after particle radiotherapy, where in-vitro experiments with particle beams provided the first evidence that increased LET and consequently non-repairable DSB may limit compensatory survival mechanism elicited by the TMs.

## Materials and Methods

### Cell culture

The primary patient derived glioblastoma stem-like cell lines (GBSC) S24 [4] and T269 [19] were cultured as 3D spheres in Dulbecco’s Modified Eagle Medium/F-12 (DMEM/F-12) with 15 mM HEPES and L-Glutamine. The medium was enriched with 5 µg/mL each of Heparin and Insulin, 1X B27, and 20 ng/mL of both Epidermal Growth Factor (EGF) and Fibroblast Growth Factor (FGF). The cultures were maintained at 37°C in a 5% CO_2_ environment. All reagents were sourced from Thermo Fisher Scientific, except for Heparin and Insulin, which were obtained from Sigma-Aldrich. Both cells lines were transduced with GFP lentiviral constructs as detailed in [3]. For monolayer cell culture, cells were seeded in Poly-L-Lysine (PLL; 1:100 in sterile water) -coated 96 well plates or 18-well chambers (Ibidi, Germany).

### Irradiation treatment

X-rays irradiation was performed using X-Rad320 cabinet irradiator (Precision X-Ray Inc, North Branford, CT) at 320 keV with a dose rate of 110 cGy/min. For protons, helium and carbon ion delivery, a 4×4×4cm^2^ spread-out Bragg peak (SOBP) was physically optimized to deliver doses of 1, 2, 4 and 6 Gy, as previously described [X]. For each experiment, airgaps within the wells in 96-well plates were filled with 3 % agarose gel to replace air with a water-equivalent material (improve target homogeneity). To reduce physical uncertainties introduced by the heterogeneous target, a detailed geometry of the utilized 96-well plates was incorporated into a FLUKA Monte Carlo (MC) simulation of the HIT beam-line [20-22].

### Clonogenic survival assay

Clonogenic survival assay was performed using S24 cells in a 96-well format as described previously [23]. To maintain 3D sphere culture and enable sphere attachment and colony formation, single cells were resuspendend in cell culture media containing Matrigel (1:100; Corning, Netherlands) and seeded on previously coated with Matrigel (1:50) 96-well plates (Ibidi, µ-plate 96). Cells were seeded as single cells and allowed to attach before irradiation. Non-irradiated cells were used as controls. Post-irradiation, cells were incubated for colonies formation (9 days) and subsequent imaging using a live cell analysis system (whole well imaging) at 4x magnification (IncuCyte, Essen Bioscience, Sartorius, Germany). Images were analyzed by the IncuCyte Zoom Software (Essen Bioscience) and colony counts were verified manually.

### Immunofluorescence staining

Cells were fixed 1 day after irradiation using 4% paraformaldehyde (PFA) for 20 minutes and blocked with 1% BSA, 0.1% Triton X in PBS for 25 minutes at room temperature (RT). γ-H2AX labeling was performed using primary anti-γ-H2AX antibody (1:100, Cell Biolabs, San Diego, CA, USA) and secondary Alexa 488-conjugated anti-mouse secondary antibody (1:600, Life Technologies, Germany). Nuclei were counterstained with 4’,6-diamidino-2-phenylindole (DAPI; 1 µg/Ml in PBS).

### Fluorescence microscopy

Images for γH2AX foci and nuclei were acquired by an Olympus IX83 widefield microscope (20X objective) and a tunable Lumencor Spectra-X LED. Samples containing colonies (S24 cells) were imaged with an Olympus IX83 widefield microscope using a 10x objective array 9 days post-irradiation, whereas cell monolayer samples (S24 and T269 cells) were imaged with an 20x objective, 12 days post-irradiation. Using the Olympus ScanR (v3.2) high throughput imaging software, the wells of interest containing the irradiated and control cells were selected, and 9 non-overlapping multi-channel micrographs per condition were captured. Each micrograph consisted of GFP signal. All images were captured using a Hamamatsu Orca Flash v2 sCMOS with a pixel size of 0.66 (10X) or 0.33 (20x) µm.

### Image processing and analysis

γ-H2AX signal was analyzed automatically using in-house developed macros for FIJI / ImageJ. In short, images of DAPI were background subtracted using Rolling Ball algorithm, blurred with Gaussian blur, segmented with Find Maxima tool and thresholded. The output of Find Maxima was Single Points (centers of DAPI stained nuclei). The images of stained area were processed with Rolling Ball Background Subtraction, Gaussian Blur and the selection was created above threshold. All single points (centers of DAPI) within the selected stained area were counted and compared to the total cell count. Single γ-H2AX foci area was analyzed using Olympus ScanR analysis package (v3.9). Rolling ball background subtraction was performed for γ-H2AX signal. Thresholding detection method was used to detect γ-H2AX and DAPI, where one γ-H2AX focus area was defined as an area of at least 2 pixels.

Colonies sample images were processed with constant settings using the Olympus ScanR analysis package. Rolling ball background subtraction was performed for GFP signal. Thresholding and edge detection method was used to detect colonies and TMs. Colonies were counted and TMs were gated out and segmented from colonies based on the elongation and circularity factors (>4 and > 1.8, respectively). TMs were then counted and for each ROI average TMs signal area was measured. To measure and quantify microtubes and count cells in a monolayer, in-house developed macro for ImageJ was used. In short, images of microtubes were pre-processed with Median filter to create selections of large structures (cell bodies). These selections were used back on original images to exclude large structures from further analysis. The images were thereafter thresholded, converted to mask, skeletonized and areas of microtubes were measured with Analyze Particles tool. To count cells, images were blurred with Gaussian blur, thresholded, cells bodies were segmented with Find Maxima tool and cells were counted with Analyze Particles tool. In both analysis cases, TMs density was normalized to the number of surviving colonies or cells. For each treatment condition, TMs density fold change was calculated as a total TMs signal area within ROI (1321.29 x 1321.29 µm each ROI, n= 9) – TMs density, and normalized to the average TMs density of respective control samples. TMs > 50 µm length were considered in analysis independently of their connectivity status to the neighbouring cells.

### Statistical analysis

Data were plotted and analysed using Python and GraphPad software. One-way ANOVA (Tukey′s multiple comparisons test, were performed with GraphPad Prism software. Data are presented as averages and 95% confidence interval error bars, unless stated differently. Data fitting and correlation analysis were done using Python module. LQ-fitting was performed on the clonogenic survival data using an in-house tool based on Minuit package available in ROOT [24] as in previous reports [25].

## Results

### X-rays induce TM formation in patient-derived GB cells

To study the effects of radiotherapy on tumor microtubes (TMs) *in vitro*, patient derived glioblastoma stem cells (GBSC) S24 and T269 were employed. Both models are paradigmatically utilized for investigating the role and implications of TMs in GB progression and therapy [2-4, 26]. X-ray irradiation induced an increase in TMs in both cell lines. This increase was dose apparent in both cell lines. S24 cells displayed high expression of TMs already with starting dose of 1 Gy, which remained high over all tested dose levels (Fig. 1a). T269 cells displayed a lower expression of TMs compared to S24 cells at the given magnification and conditions. Nevertheless, it was found that X-ray irradiation could enhance TMs, when doses up to 4 Gy were applied (Fig. 1b). However, when the dose was increased to 6 and 8 Gy, there was a decrease in TMs density, a contrast to the response observed in S24 cells. Consistent with the fact that S24 GBSCs were obtained from a relapsed and thus tendentially more resistant GB, whereas T269 was derived from a newly diagnosed GB (19), clonogenic survival analysis indicated higher resistance of S24 cells post X-rays exposure compared to the T269 and other GB cell lines [27, 28]. Demonstrating this, the S24 surviving fraction at 2 Gy SF_2_= 0.73 ± 0.16 (Fig. 1c), confirming higher-end resistance compared to the T269 (SF_2_= 0.26 ± 0.04), while cell killing was dose dependent in both. Further analysis revealed that survival of both cell types correlated to the TMs density change in dose dependent manner. However, this correlation was higher in more sensitive T269 cells (Fig. 1d).

**Figure 1.**
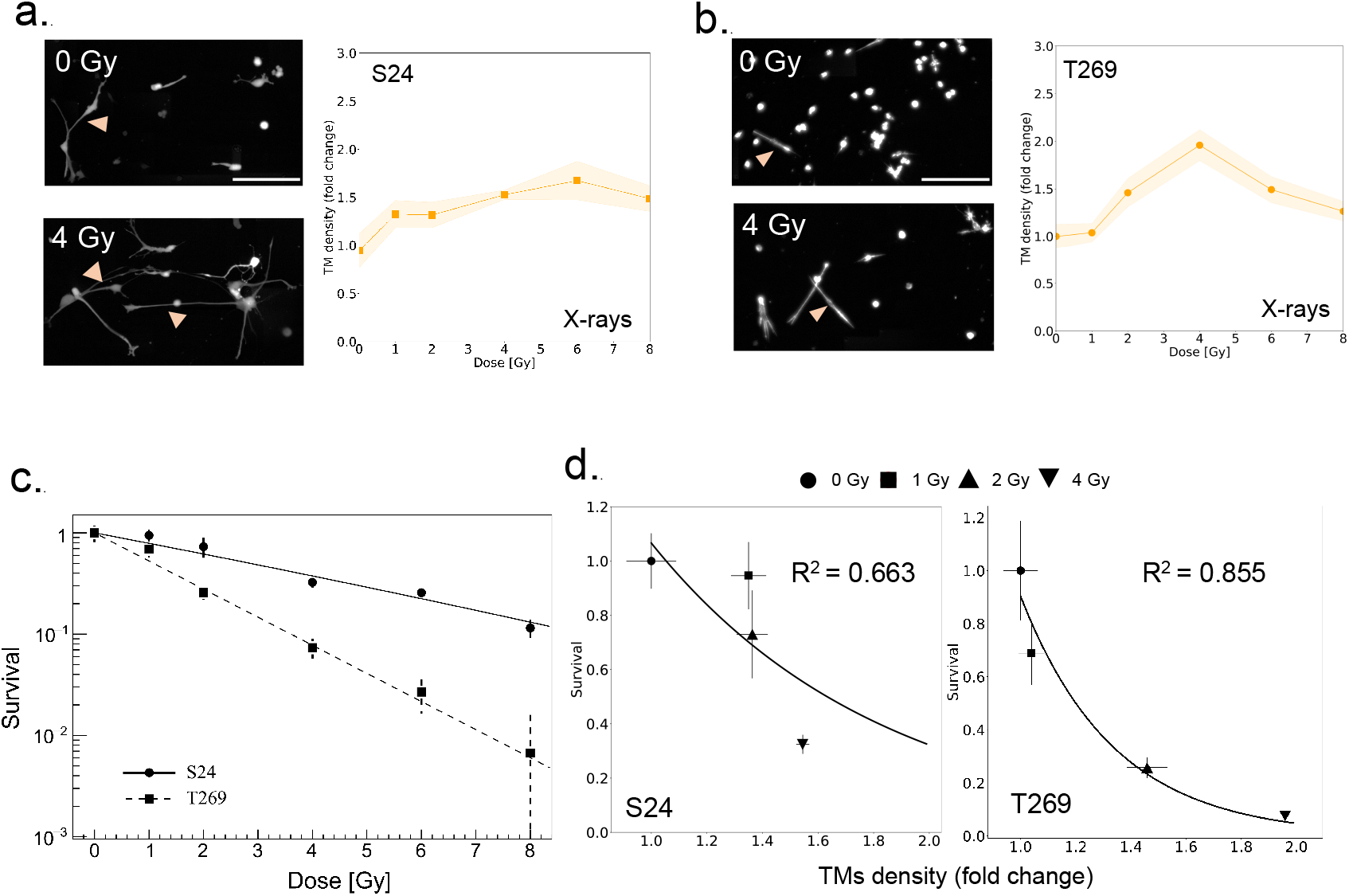
GB response to radiotherapy and TM formation. Representative images and quantification of TMs density in **a.** S24 and **b**. T269 GB cells. Data are presented as means and connecting lines with 95% confidence interval (CI, n= 6). **c**. Clonogenic survival of S24 and T24 cells after X-rays irradiation. Data points represent means and standard errors (n= 6). Line represents linear quadratic (LQ) fit, describing survival curve behavior. **d**. Correlation of GB cell survival and TMs density. Data are means and standard errors.

### Particle irradiation with higher LET limits TMs density

We hypothesized that increased LET, causing more unrepairable DNA damage could lead to decrease in TMs network density (Fig. 2a). To address this, the effect of clinically relevant LETs for tumor targeting at the different dose and for different particle modalities on TMs networks was investigated in S24 cells (Fig. 2b). Proton irradiation (^1^H, 6 keV/µm) induced a partially dose-dependent increase in TMs density, peaking at 4 Gy. Helium ions (^4^He 35 keV/µm) also increased TMs network density, albeit to a lesser extent than protons. In contrast, higher-LET carbon ion treatment (^12^C 107 keV/µm) TMs density remained similar to the one in control sample (0 Gy), across all administered doses (Fig. 2c). DNA damage investigation revealed modality e.g. LET-dependent increase in DNA damage. This was evidenced by a rise in the number of RIF-positive cells and the RIF area following 2 Gy irradiation, observed on the first day post-treatment. The effect peaked after carbon irradiation (Fig. 2d). LET dependent TMs network decrease for the same dose (Fig. 2e) highly correlated with increase in unrepaired RIF area (R^2^= 0.86, Fig. 2e), serving as an indicator of complexity of DNA damage [16, 29]. In addition to S24 cells, LET-dependent TM decrease was also apparent for T269 cells (Fig. 2f).

**Figure 2.**
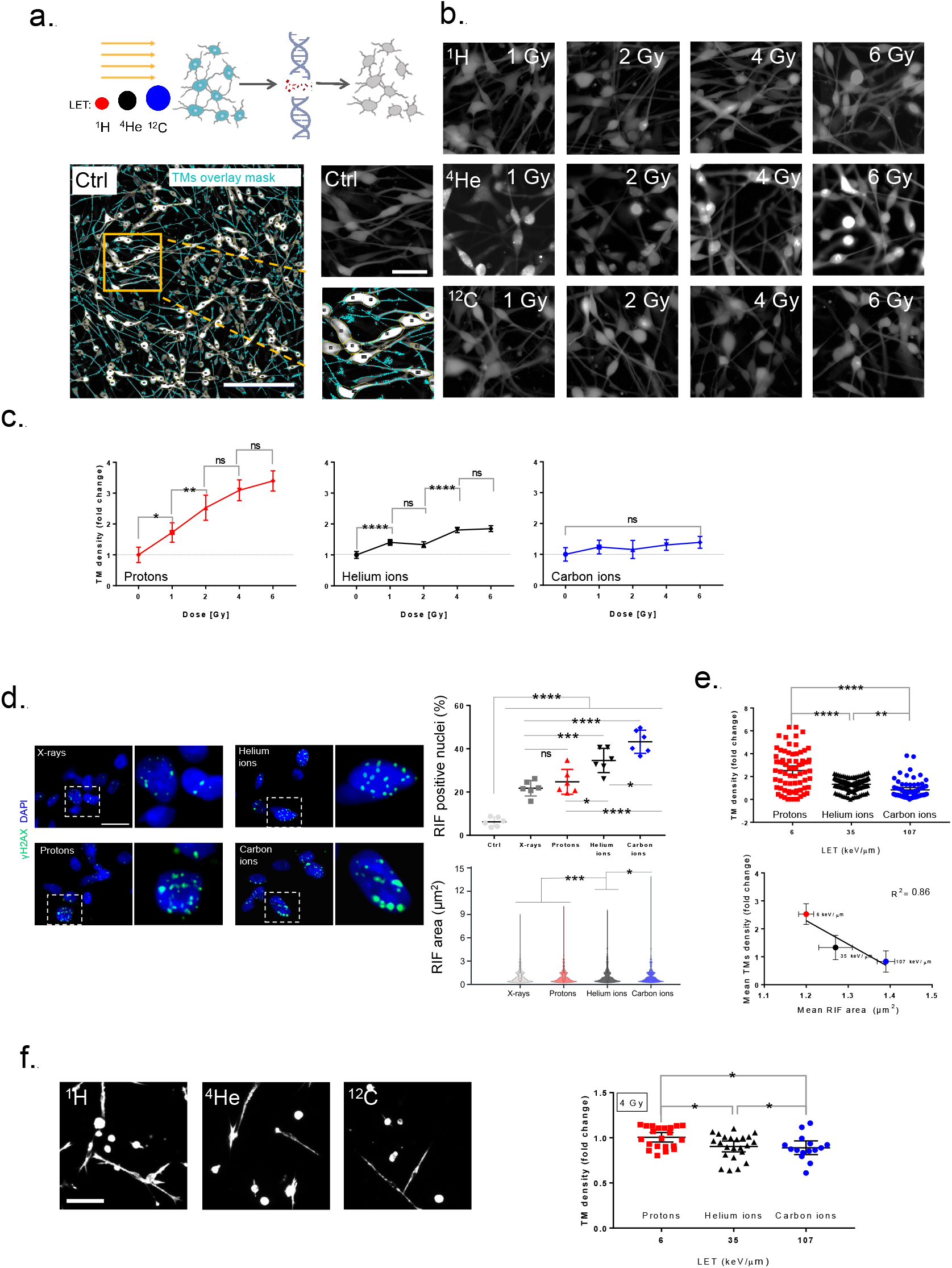
TMs destabilization as a function of particle irradiation modality. **a.** Schematic of TMs network destabilization post-particle irradiation. **b**. Overlay image for a control sample indicates TMs segmentation using ImageJ′s macro. Scale bar: 400 µm (blue lines: TMs mask, yellow lines: segmented cell bodies) and representative images of TMs network in S24 cells after irradiation with protons (^1^H), helium (^4^He)- and carbon ions (^12^C) at different doses. (LET: 6, 35, 107 keV/µm, respectively). Scale bar: 100 µm. **c**. TMs density fold change after protons, helium- and carbon ions irradiation in S24 cells. Data are normalized to the control (dashed line; 1) and are presented as means and error bars (95% CI, n= 6; One-way ANOVA with Tukey’s comparison; **** p< 0.0001, ** p< 0.01, *p< 0.05). **d**. DNA damage γH2AX foci in S24 cells 1 day after 2 Gy treatment using different radiation qualities (scale bar= 25 µm, n= 6 samples, each sample contained > 1000 nuclei). Data are means and standard errors compared using one-way ANOVA **** p< 0.0001, *** p< 0.001, *p< 0.05). **e**. Comparison of the TMs density fold change after irradiation with different particles at 2 Gy dose (top panel, n= 6, One-way ANOVA with Tukey’s comparison; **** p< 0.0001) and correlation of radiation induced γH2AX foci (RIF) area to TMs density as a function of LET at 2 Gy (bottom panel ; data are means and standard errors). **f**. Comparison of the TMs density fold change in T269 cells after irradiation with different particles at 4 Gy dose (panel left, n= 6, One-way ANOVA with Tukey’s comparison; 95% CI, *p< 0.05)

### Particle radiotherapy decreases cell survival as a function of dose and LET

Choosing an optimal LET for irradiation can be achieved not only by applying different particle modality irradiation, but could be further optimized for each particle type. In the context of the more resistant S24 cells, we present the survival rates as a function of dose across LETs using protons (Fig. 3a), helium ions (Fig. 3b) and carbon ions (Fig. 3c). The data reveal an increased potential for cell eradication corresponding to an increase in both LET and dose across all three irradiation modalities. The most pronounced LET, and consequently, the highest cell killing was observed after carbon ion irradiation at 6 Gy dose, resulting in a survival fraction of ∼0.08 ± 0.004. This is in contrast to the survival rates of ∼0.26 ± 0.02 for X-rays, ∼0.25 ± 0.02 for protons, and ∼0.19 ± 0.07 for helium ions, underscoring the elevated cell-killing potential of high-LET carbon ions.

**Figure 3.**
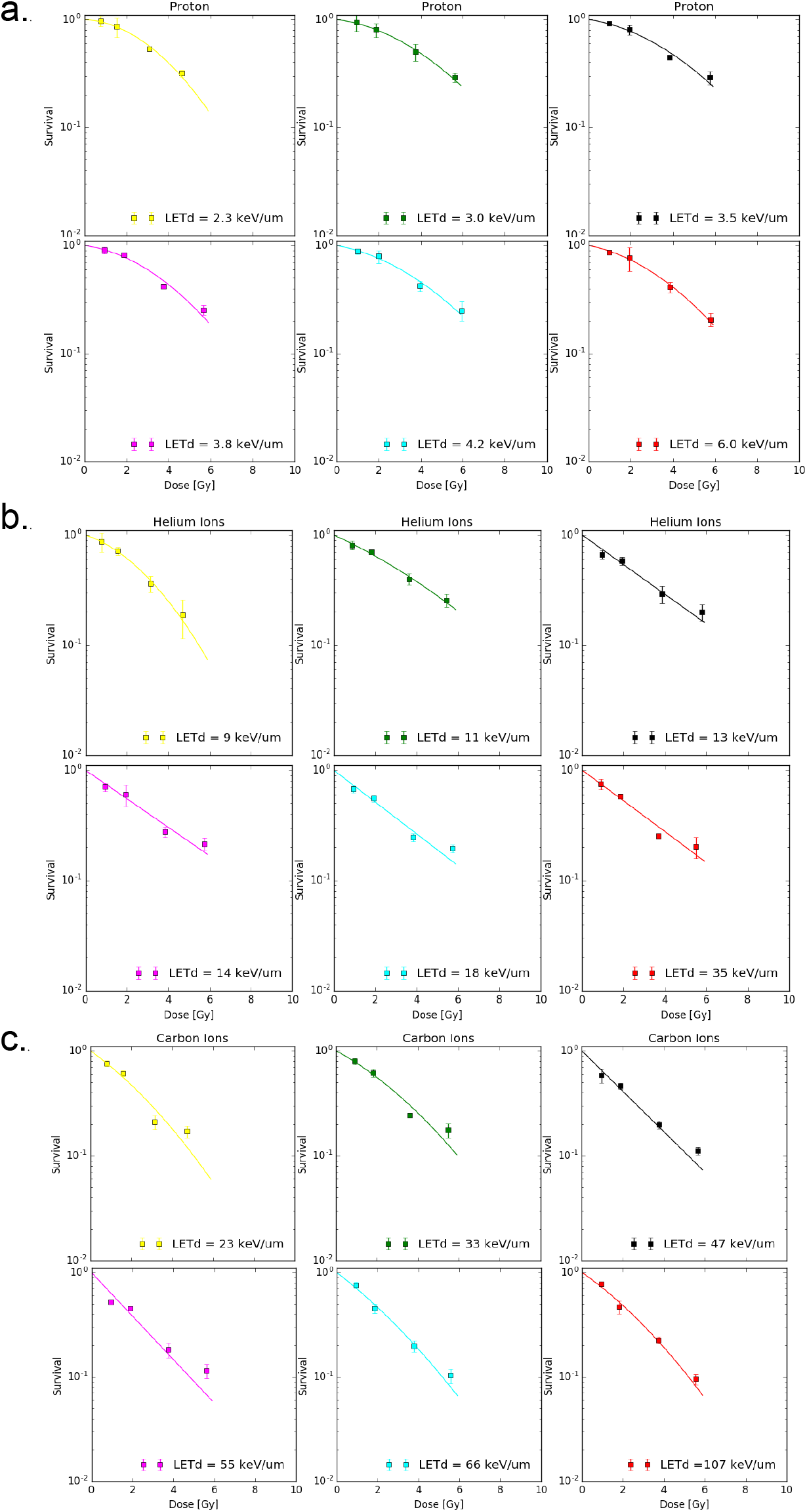
GB cell survival decreases with increase in dose and LET. Clonogenic survival of S24 cells after irradiation with **a.** protons, **b**. helium ions, and **c**. carbon ions as a function of dose and LET. The points with standard error bars represent the experimental data (n= 3); LQ fits are depicted with solid lines.

To understand the impact of varying LET of each particle irradiation on TMs, we investigated TMs formed not only among single cells (Fig. 1 and Fig. 2), but also among surviving cellular colonies. Proton irradiated samples revealed TMs connecting S24 colonies, whereas this was less prominent in helium and carbon irradiated samples (Fig. 4a). In carbon ion irradiated samples, most of the colonies exhibited disconnected TMs (Fig. 4a). Similarly, to X-rays (Fig. 1) low-LET beams, particularly protons (Fig. 4b), and to lower extent helium ions (Fig. 4c), led to an increase in TMs for doses > 1 Gy. Maximal LETs were approximately 6 keV/µm for protons and 18 for helium ions. Likewise, low-LET carbon ions (33 and 66 keV/µm) irradiation resulted in a moderate increase in TMs for irradiations < 6 Gy. Upon escalating carbon ions LET to clinically relevant value (107 keV/µM), the TMs density was reduced to the one of a control for all tested doses (Fig. 4d).

**Figure 4.**
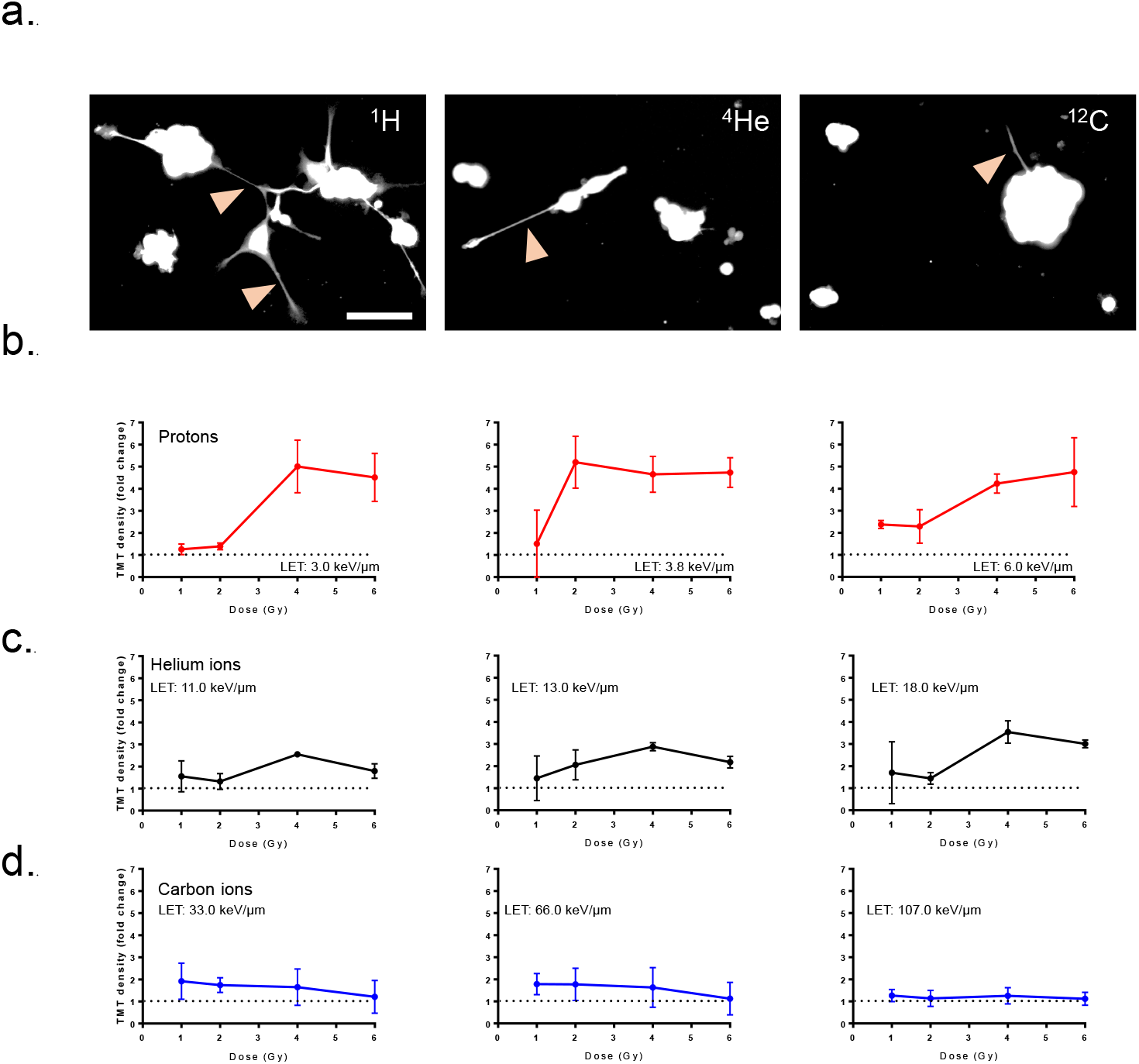
GB colonies cross-connect with TMs. **a.** Representative images for S24 inter-colony TMs (arrow) after 6 Gy irradiation with protons (LET: 3 keV/µm), helium (LET: 13 keV/µm)- and carbon ions (LET: 66 keV/µm). Scale bar: 100 µm. Inter-colony TM density fold change using different LETs for **b**. protons, **c**. helium ions and **d**. carbon ions. Dashed line represents TMs density in control sample and it is normalized to 1. Data are presented as means and standard errors.

## Discussion

TMs are elongated cytoskeletal-enriched membrane tubes that connect tumor cells into functional networks. They play a significant role in the progression of GB, a highly aggressive brain tumor, by facilitating diffuse invasion and therapy resistance [30]. Studies have shown that TMs contribute to the resistance of GB cells to standard radiotherapy with X-rays [3, 4]. Cells connected by TMs exhibit prolonged viability post-X-ray treatment and an increase in TMs following irradiation [4]. The comparison between the TM-richer and more resistant GB model (S24) and the TM-poorer and more sensitive GB model (T269) revealed distinct differences in their responses to X-ray irradiation. In T269 cells, TM density could be altered at doses exceeding 4 Gy, indicating that the impact of irradiation on TM formation may vary among different GB cells, potentially influenced by their connectivity signatures [26] or DNA damage response. Since with the dose increase DNA complexity increases [31], this may suggest a potential link between the complexity of DNA damage, residual (unrepaired) DNA damage foci (radiation induced foci; RIF), and the downregulation of TM networks. Induction of complex damage could be also achieved with lower irradiation doses using high LET irradiation [31, 32]. This type of damage is harder for cancer cells to repair [17], making high LET irradiation particularly effective in targeting radio-resistant tumors.

To this point, there is no data on the response of GB TM networks to particle irradiation therapy. Each type of irradiation particle has unique benefits and drawbacks, highlighting the need for comprehensive biological investigations across all particle types at varying LETs for optimal clinical use. The DNA damage response varies with different radiation qualities, reflecting their distinct modes of action. Low linear energy transfer (LET) radiation, such as X-rays or gamma rays, primarily induces indirect DNA damage through radiolysis [33]. In contrast, high-LET particles directly cause complex double-strand break damage [34], also demonstrated in this work for GB cells, suggesting that low- and high-LET radiation exhibit different DNA damage processing kinetics [14, 16, 35].

Our findings present the first evidence that high-LET particle beams significantly impact GB cells survival and the structural integrity of TM networks, correlating with non-repairable DNA damage. Even though low-LET X-rays irradiation led to a different dose response of TM-poor vs. TM-rich GB cells, high-LET carbon irradiation resulted in a similar outcome for cell types. This aligns with pre-clinical studies that highlight the efficacy of carbon ions in eradicating GB tumor models [28, 36, 37]. This effectiveness is attributed to unique potential of carbon ions to induce complex-unrepairable DNA damage [16, 38], elicit a distinct molecular response [39] and exhibit a reduced reliance on oxygen presence and intracellular oxidative stress formation compared to X-rays [36].

## Conclusion

The findings presented here provide valuable insights into the effects of particle irradiation on GB cells, particularly in relation to TMs formation, and highlight the potential of particle radiotherapy as a therapeutic strategy for GB. This work represents the ground work in utilizing high-LET particle irradiation for disconnecting TMs and underscores the need for preclinical in vivo research to fully understand the implications of these treatments on TMs dynamics and GB progression.

## Acknowledgments

The authors express their gratitude to Prof. Wolfgang Wick and Prof. Frank Winkler for valuable discussions and insightful input that contributed to this manuscript. The authors thank Ms. Nora Schuhmacher and Claudia Rittmüller for their excellent technical support.

